# Virally encoded interleukin-6 (vIL-6) coordinates with human IL-6 to modulate cytokine expression during KSHV infection

**DOI:** 10.1101/2025.10.02.679950

**Authors:** Yahaira Bermudez, Samantha Schultz, Jacob Miles, Will Dwyer, Mandy Muller

**Affiliations:** Department of Microbiology, University of Massachusetts, Amherst, USA; Molecular and Cellular Biology program, University of Massachusetts, Amherst, USA; Institute of Microbiology, University Hospital of Lausanne, University of Lausanne, 1011 Lausanne, Switzerland

## Abstract

Kaposi Sarcoma-Associated Herpesvirus (KSHV) is characterized by its ability to establish lifelong infections that can lead to disease in immunocompromised individuals. Understanding the complex interactions between KSHV, the host cellular machinery, and the immune response is essential for unraveling the mechanisms underlying viral persistence and pathogenesis. KSHV extensively remodels the host gene expression landscape by inducing widespread RNA decay *via* the viral endoribonuclease SOX. While the vast majority of cellular mRNAs are degraded by SOX, select transcripts including Interleukin 6 (IL-6) escape degradation. Interestingly, KSHV encodes a viral homolog of IL-6, known as viral interleukin-6 (vIL-6). Given the importance of IL-6 for KSHV biology and its ability to escape SOX-mediated decay, we explored how vIL-6 expression is regulated upon KSHV infection. Our data demonstrate that unlike IL-6, vIL-6 3’UTR does not provide protection from SOX degradation, yet vIL-6 mRNA steady state levels remain high throughout KSHV lytic cycle, suggesting a distinct transcriptional regulation. To assess the contributions of vIL-6 and IL-6 in modulating host gene expression, inflammatory signaling and KSHV-driven pathogenesis, we used a vIL-6 null viral mutant combined with an IL-6 knock down to perform transcriptomic analyses. This analysis revealed that depletion of IL-6, vIL-6 or both results in distinct alterations of the host gene expression environment, with differential effects on chemokine signaling and related pathways as well as a synergistic effect when both IL-6 and vIL-6 are present. Furthermore, functional assays confirmed vIL-6 alone is sufficient to activate key signaling pathways and pro-inflammatory responses, including the JAK/STAT pathway. Collectively, our findings suggest a mechanistic synergy between vIL-6 and IL-6 that may enhance KSHV-mediated immune modulation and contribute to viral persistence and tumorigenesis.

## Introduction

It is estimated that 15% of all human cancers are linked to oncogenic viruses, representing a significant burden for human health worldwide^1^. Kaposi’s sarcoma-associated herpesvirus (KSHV), a gamma-herpesvirus is associated with the development of several neoplasms afflicting immunocompromised individuals. Among these is its namesake disease Kaposi’s sarcoma (KS), as well as primary effusion lymphoma (PEL) and multicentric Castleman’s disease (MCD)^2^. KS in particular, has been a major focus of research, as it is one of the most common and aggressive cancers in untreated AIDS patients^3,4^. KSHV life cycle alternates between two distinct stages: a long-lasting latent phase and a productive lytic phase^5^. Like other herpesviruses, KSHV infection persists for the lifetime of its host and is maintained for decades in the latent phase. On the contrary, the lytic phase is tightly controlled and often suppressed to minimize encounters with the host’s immune system. However, in KS lesions, the lytic cycle is instead not repressed and plays a decisive role in the pathogenesis of the tumors. KSHV lytic factors are critical for tumor formation and maintenance, likely due to secretion of paracrine factors that promote tumor development^2^. Therefore, deciphering the regulation of this lytic phase is pivotal to make progress on our global understanding of KSHV-associated neoplasms.

During the lytic phase, KSHV induces a striking reprogramming of host gene expression known as *Host Shutoff*. A hallmark of this process is the broad degradation of cytoplasmic mRNA, which is believed to suppress immune activation and divert cellular resources toward viral replication^6-8^. Without this widespread mRNA clearance, KSHV exhibits major impairments in its lifecycle *in vivo* which critically disrupts its ability to cause disease and drive cancer development^6,8^. This global mRNA decay is orchestrated by a single viral protein, SOX. While SOX is broadly engaged in degrading host transcripts, we have identified a subset of host genes that are able to evade this viral silencing^9-12^. Notably, although SOX targets the majority of the host transcriptome, some host mRNAs remain resistant to cleavage, even when they contain strong SOX recognition motifs. The best characterized SOX “escapee” is IL-6, a pro-inflammatory cytokine that has been shown to be critical to maintain KS tumors and promote angiogenesis of the KSHV-associated malignancies and diseases^9,11,13^. Intriguingly, KSHV also encodes its own viral mimic of IL-6: the vIL-6 protein encoded by the K2 gene.

Recent studies on vIL-6 have demonstrated that it is highly expressed in KSHV replicating cells while also expressed in a small population of latently infected cells^14,15^ and all KSHV-associated malignancies have detectable vIL-6 levels^16,17^. vIL-6 contributes to enhancing cell proliferation, cell migration, and angiogenesis^18^. In mice, vIL-6 also supports tumor metastasis in murine xenograft models^19^. vIL-6 is broadly described as binding the IL-6 receptor (IL-6R). However, IL-6 has an absolute requirement for both the IL-6RΔ and the gp130 subunit, while vIL-6 appears to solely rely on the transmembrane gp130 subunit^20^. Once the receptor is engaged, similarly to IL-6, vIL-6 can activate the JAK/STAT pathway^21^. In addition, vIL-6 activates the AKT pathway promoting some key oncogenic phenotypes^22,23^ as well as VEGF expression resulting in increased angiogenesis^24^. One major difference between IL-6 and vIL-6 is that IL-6 is rapidly secreted from cells while vIL-6 is retained primarily within the endoplasmic reticulum (ER)^25^. Interestingly, IL-6 may play a role in enhancing the growth and tumorigenicity of B cells immortalized by another herpesvirus, EBV (Epstein Barr Virus)^26^. Given the known role of human IL-6 in inflammation^27^ and the roles of vIL-6 in KSHV associated tumorigenesis, we set out to understand their regulation and interplay during infection.

Here, we show that unlike IL-6, the vIL-6 transcript is not directly refractory to SOX. However, its expression increases during lytic infection, suggesting differential transcriptional regulation between the two transcripts. Knocking down IL-6 has a more pronounced effect on host gene expression than a vIL-6-null mutation during lytic reactivation, while IL-6 silencing in a vIL-6 null background leads to combined modulation of inflammatory signaling pathways. Taken together, these findings provide important insights into the complex regulatory network driving inflammatory signals during KSHV lytic reactivation.

## Results

### Comparative expression of vIL-6 and hIL-6 during KSHV lytic reactivation

We and others have demonstrated that the pro-inflammatory cytokine IL-6 stringently resists SOX-mediated RNA decay. This resistance stems from an RNA stability element located within IL-6 3’UTR, referred to as the <u>S</u>OX <u>R</u>esistance <u>E</u>lement (SRE). To evaluate whether the KSHV-encoded IL-6 mimic, vIL-6, similarly contains a putative SRE-like element in its 3’UTR, we designed a GFP reporter (**Fig 1A**) as previously used^10-12^. The vIL-6 transcript is encoded on the reverse strand of the KSHV genome and its 3’UTR is ∼100 nucleotides long^28^. We cloned the vIL-6 3’UTR downstream of GFP (vIL-6-3’UTR) and found that, contrary to the IL-6 3’UTR, it was unable to confer protection from SOX in transfected HEK-293T cells (**Fig. 1A**). This was surprising as vIL-6 is known to be highly expressed throughout KSHV infection and in particular, during the lytic cycle when SOX is active. We thus next compared vIL-6 RNA steady state levels to IL-6 levels in the KSHV positive cell line iSLK.219 and the PEL-derived TREX-BCBL1s. As shown in **Figure 1B and 1C**, IL-6 mRNA slowly increases over the lytic cycle demonstrating its ability to escape SOX decay as expected. Given our above results that vIL-6 seems to be susceptible to SOX, we were surprised to observe that vIL-6 mRNA increased exponentially over the KSHV lytic cycle. We thus next wanted to evaluate whether vIL-6 protein levels would similarly dramatically increase during the lytic cycle. Intriguingly, vIL-6 protein levels remained stable throughout lytic reactivation (**Fig. 1D**). Given that IL-6 is primarily secreted, we next performed an ELISA to quantify IL-6 protein levels in the cells and in supernatants. In iSLK.219 cells, we observed that IL-6 protein expression increases and remains stably secreted throughout the KSHV lytic cycle (**Fig. 1E**). However, in BCBL-1 cells, we barely detected IL-6 intracellularly but IL-6 was quickly secreted (**Fig. 1F**). Taken together, this suggested that perhaps vIL-6 transcription was significantly activated during lytic reactivation as a means to compensate for SOX decay and thus maintaining a constant supply of translationally competent vIL-6 mRNA for vIL-6 protein production. To monitor whether transcription of IL-6 and vIL-6 is differentially regulated, we pulse-labeled cells with 4-thiouridine (4sU) for 10 min. 4sU is incorporated into actively transcribing RNA and can be subsequently coupled to HPDP-biotin and purified over streptavidin beads, then quantified by RT-qPCR to measure nascent transcript levels^29^. 4sU-labeled RNA levels were also normalized to 18S rRNA, which is produced at a constant level in the presence and absence of SOX. We observed that both IL-6 and vIL-6 undergo a rapid burst of expression, but as opposed to IL-6, vIL-6 exponentially increases over the lytic cycle (**Fig.1G**). This could suggest that transcription rates of both vIL-6 and IL-6 are complementary during infection: IL-6 achieves high-level expression by resisting SOX-mediated RNA decay and therefore is less demanding transcriptionally, while vIL-6 compensates for RNA degradation through increasing transcriptional activation.

**Figure 1.**
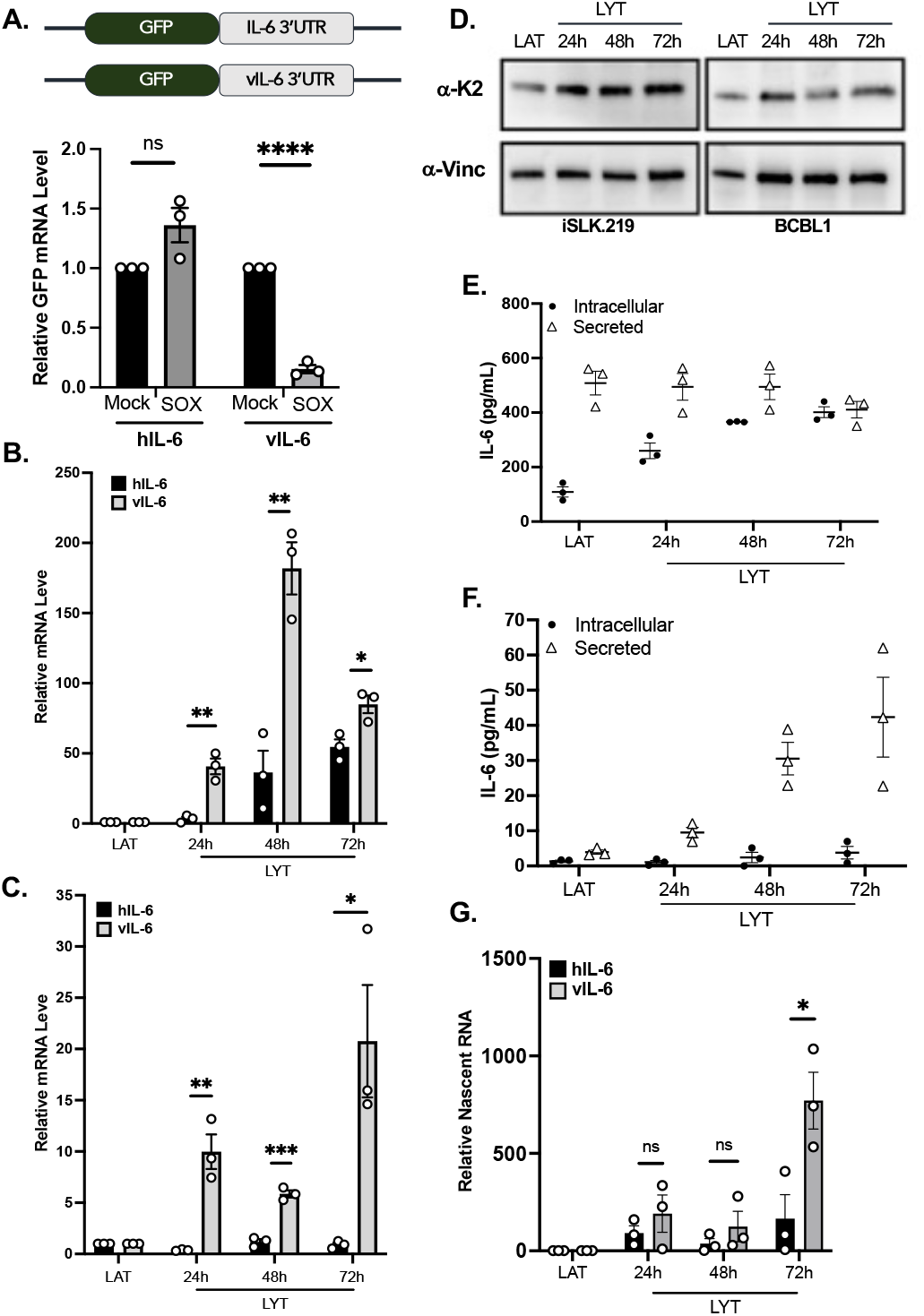
vIL-6 does not resist SOX but remains stable during KSHV lytic infection. **(A)** HEK293T cells were transfected with the indicated GFP reporter (top) containing either the 3’UTR sequence of IL-6 or vIL-6 along with a control empty vector (mock) or a plasmid expressing SOX. After 24h, total RNA was harvested and subjected to RT-qPCR to measure GFP mRNA levels. **(B-C)** iSLK.219 (B) or TREX-BCBL1 **(C)** were either left latent (LAT) or reactivated (LYT) and total RNA was harvested at the indicated timepoints and subjected to RT-qPCR to measure IL-6 and vIL-6 steady state levels. Similarly, vIL-6 protein levels were measured by western blot with Vinculin used as a loading control (D) and IL-6 levels were measured by ELISA in ISLK.219 cells (E) and BCBL1 (F). (G) RT-qPCR quantification of 4sU-labeled RNA at the indicated times. Fold changes were calculated from Ct values normalized to 18S rRNA in reference to latent cells. All graphs here and after display mean ± SEM with individual biological replicates represented as dots. Statistically was determined using student *t* tests *p<0.05, **p<0.01, ***p<0.001.

### Transcriptomics analysis reveals common targets of IL-6 and vIL-6

Given that IL-6 and vIL-6 have different kinetic of expression but are known to modulate similar pathways, we next wondered whether we could isolate the contribution of IL-6 *vs*. vIL-6 at a larger scale. To do so, we took advantage of a novel KSHV vIL-6-null mutant^30^. In addition, to assess the contribution of IL-6 in these infected cells, we knocked down IL-6 expression using RNAi. Cells infected with the KSHV vIL-6 null (Δ-vIL-6) or the revertant control were then treated with siRNA targeting IL-6 (or Scramble control) and left latent or reactivated for 48 hours. Total RNA was extracted, polyA selected and used to perform RNA sequencing (**Figure 2**). Firstly, we performed hierarchical clustering on viral gene expression. Unsurprisingly, we observed that expression was primarily clustered based on Lytic vs. Latent samples. However, we noticed that knocking down IL-6 had little effect on viral gene expression: only ORF60 and ORF49 seemed to be repressed by IL-6 expression as we observed a 10-fold increase in expression in cells lacking IL-6 expression. Clustering analysis also revealed that our expression profiles segregated based on vIL-6 expression, as several viral genes were no longer detectable when vIL-6 was not expressed. For host genes (**Fig 2B**), as expected, we observed that as opposed to viral gene regulation, IL-6 had more of an impact on gene expression rather than vIL-6, with many genes involved in chemokine expression affected by IL-6 knock down. Interestingly, when both IL-6 and vIL-6 are knocked down, we observed a synergy in expression modulation, suggesting that IL-6 and vIL-6 work together to modulate these pathways.

**Figure 2.**
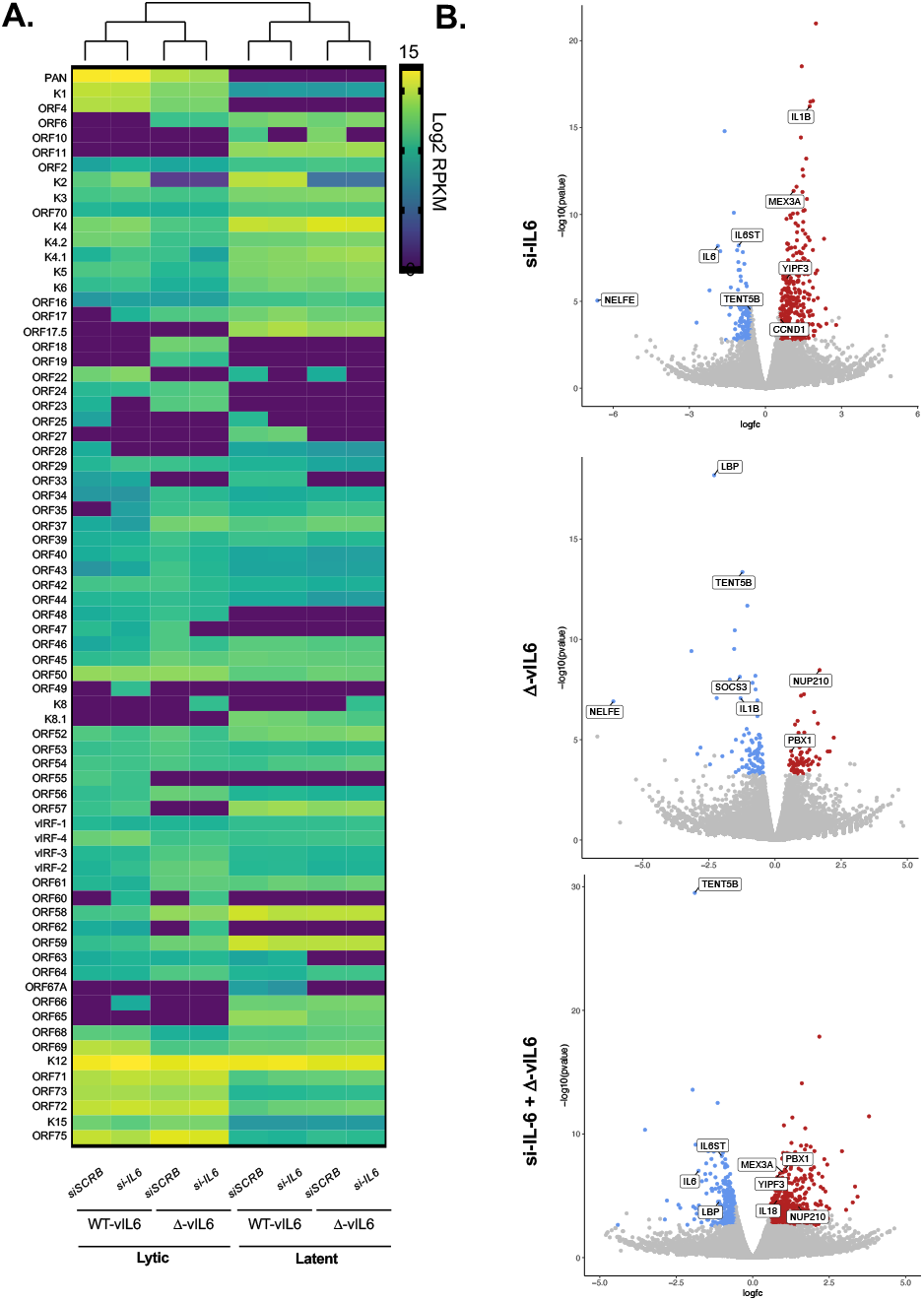
IL-6 and vIL-6 coordinate together to control KSHV chemokine patterns. Cells were infected with a WT KSHV virus or vIL-6 null mutant into cells pre-treated with an siRNA targeting IL-6 (or Scramble-SCRB control). Cells were then left latent or reactivated for KSHV to enter its lytic cycle. Total RNA was harvested and used for RNAseq. Viral genes (A) expression was measured and the heatmap represents the relative expression. Volcano Plots (B) represent the fold change of host gene expression in the indicated conditions (over their matching controls).

After sorting and processing the data, we identified several genes with distinct expression patterns upon knock down of IL-6 and/or vIL-6. To validate these genes expression patterns, we performed RT-qPCR (**Supplementary Fig 1**). Specifically, we assessed gene expression in Δ-vIL-6 *vs*. vIL-6-REV (WT) cells treated with siRNA targeting IL-6 or scramble control, under both latent or reactivated conditions for 48 hours as above. We first examine expression of IL6ST, which encodes the gp130 receptor and which was identified by our transcriptomic approach as significantly downregulated when both vIL-6 and IL-6 expression was repressed. We also examined CCND1, a key regulator of the cell cycle, that also plays roles as a transcription factor regulator, chromatin remodeling, cellular metabolism and differentiation. This transcript appeared in our transcriptomic data as being specifically upregulated upon IL-6 knock down. Both IL6ST and CCND1 expression patterns were recapitulated by RT-qPCR (**Supplementary Fig 1**). SOCS3, a known negative regulator of the JAK/STAT pathway was selected as a validation target for vIL-6 effect. SOCS3 was found to be downregulated in our RNA-seq data, a result confirmed by RT-qPCR (**Supplementary Fig 1**).

### Induction of the JAK/STAT pathway during KSHV lytic reactivation

Human IL-6 functions are mediated through its binding to the IL-6 receptor (IL-6R) and signaling initiated by the receptor complex of IL-6/IL-6R and gp130^31-33^. Signaling from activated gp130 is transmitted by the Janus kinase (JAK)-signal transducer, the activator of transcription (STAT) axis and by the Ras-mitogen-activated protein kinase (MAPK) cascade among other cross talked pathways^21^. Interestingly, unlike hIL-6, which initially binds to the IL-6 receptor (IL-6R) and then associates with the signal-transducing receptor subunit gp130, vIL-6 has been shown to directly and indirectly bind and stimulate gp130 without requiring the IL-6R ^16,21^. As a result, vIL-6 can activate a broader range of target cells throughout the body, as all cells express gp130. Consistent with this, our transcriptomic analysis revealed differential expression of host factors involved in pro-inflammatory signaling. Therefore, we examined the contribution of vIL-6 to the induction of the JAK/STAT pathway, as well as the synergistic effect with IL-6. To investigate this, we used an ISRE luciferase reporter assay. Specifically, we transfected HEK293T cells which actively express IL6-R and gp130, with a vIL-6 expressing construct alongside the ISRE-Luciferase reporter. As expected, vIL-6 was sufficient to induce the JAK/STAT pathway compared to our control (**Fig. 3A**). Next, we measured the expression of pro-inflammatory factors upon vIL-6 expression by RT-qPCR based on candidates selected from our RNA-seq data. We observed that STAT3 mRNA expression levels increased following vIL-6 expression. (**Fig 3B**). We also observed an induction of STAT3 at the protein level upon vIL-6 expression (Fig. 3C). Moreover, another possible route through which vIL-6 and IL-6 may promote inflammation and tumorigenesis during KSHV lytic reactivation is via the activation of NF-κB^34-36^. We therefore examined NF-κB expression levels and found that, unlike STAT3, no significant increase was observed (**Fig. 3B**). To assess activation of these pathways in the context of infection, we performed ISRE-Luciferase assays and RT-qPCR assays in the KSHV infected iSLK.219 cell line. As observed in Fig. 3C, the JAK/STAT pathway was highly induced at 24 hours when host shutoff is at its peak, as viral replication ramps up and viral gene expression takes over host functions ^7,9,37^. This activation then remains stable, albeit at lower level throughout lytic reactivation (**Fig. 3D**). We next examined the expression of pro-inflammatory factors by RT-qPCR in these infected cells. As shown in **Fig 3D**, similar to HEK-293T cells, STAT3 expression increased over time during lytic infection but declined at 72 hours, while NF-κB was strongly repressed (**Fig. 3D**). Moreover, STAT3 mRNA expression levels remained constant during lytic reactivation (**Fig. 3E**). While vlL-6 alone is sufficient to activate the JAK/STAT pathway, we sought to further investigate how the absence of host IL-6 influences this signaling cascade and its downstream targets. To address this, we measured expression of vIL-6 and key pro-inflammatory factors in iSLK.219 cells treated with siRNA targeting IL-6 (**Supplementary Fig. 2**). Our results indicate that IL-6 knockdown leads to increased levels of vIL-6 supporting a feedback loop in the expression of these factors (**Supplementary Fig. 2A**). Additionally, IL-6 knockdown led to increased expression of genes associated with the JAK/STAT pathway and inflammatory signaling, such as *IL6ST, STAT3, and NF-kB* (**Supplementary Fig. 2B**). Overall, these results suggest that vIL-6 directly contributes to JAK/STAT pathway activation and the modulation of host pro-inflammatory factors during KSHV lytic reactivation, highlighting a potential mechanistic link by which synergistic signaling between vIL-6 and IL-6 promotes KSHV-associated malignancies.

**Figure 3.**
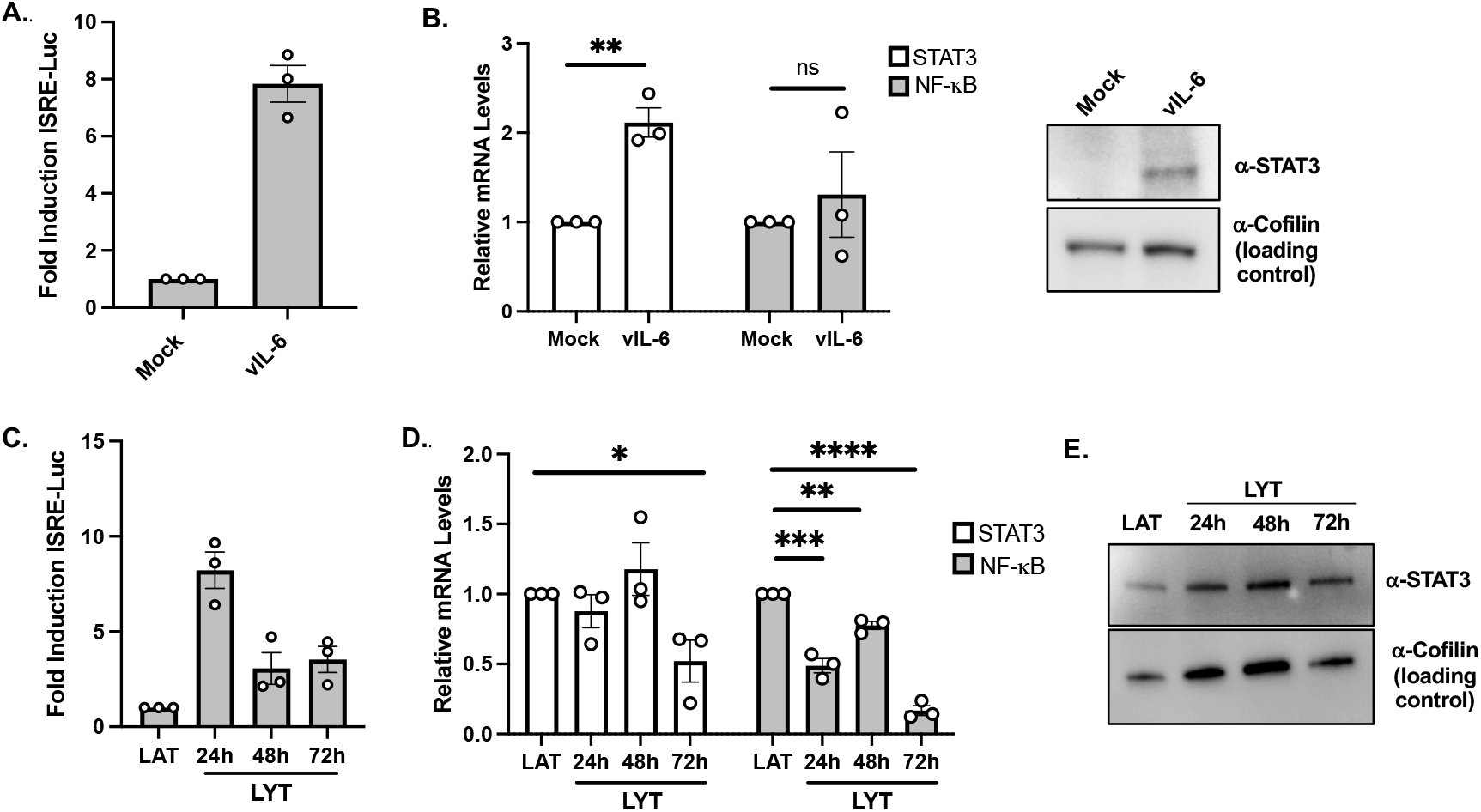
vIL-6 and IL-6 induce the JAK/STAT pathway. **(A)** HEK293T cells were transfected with a vIL-6 expressing plasmid (or mock) and the ISRE Luciferase reporter. 24h later luminescence was measured on a Berthold luminometer. (B) HEK293T cells were transfected with a vIL-6 expressing plasmid (or mock) and 24h later, RNA was extracted and used to measure STAT3 or NF-kB RNA levels by RT-qPCR (left). Protein lysates were also collected to measure STAT3 protein levels (right). (C) iSLK.219 cells were transfected with the ISRE Luciferase reporter and collected at the indicated timepoints and luminescence was measured on a Berthold luminometer. (D) iSLK.219 cells left latent (LAT) or reactivated (LYT) for 24, 48 and 72hrs were collected, total RNA was extracted and subjected to RT-qPCR to measure STAT3 and NF-κB expression levels. (E) iSLK.219 cells left latent (LAT) or reactivated (LYT) for 24, 48 and 72 hours. Cells were harvested, lysed, resolved on SDS-PAGE, and Western blotted with the indicated antibodies. All graphs represent the mean ± SEM with individual biological replicates represented as dots. Statistical significance was determined using student *t* test., ns., not significant, * p<0.05, ** p<0.01, *** p<0.001.

## Discussion

KSHV’s ability to establish persistent infections and evade immune detection, plays a crucial role in its pathogenesis and tumorigenesis. Various factors, such as host genes and virally encoded homologs of cellular genes, can influence infection progression and tumor development. One such factor, among other KSHV-encoded genes, is the viral homolog of Interleukin-6 (vIL-6)^38^. Previous studies have shown that vIL-6 shares only 25% amino acid similarity with IL-6, yet both exhibit similar functions and interactions^39,40^. Intriguingly, some studies suggest a potential antiviral function for hIL-6^41^, while others have demonstrated pro-viral effects^41,42^. Elevated expression levels of both vIL-6 and hIL-6 have been observed in the serum of patients with Kaposi Sarcoma, Primary Effusion Lymphoma (PEL), and Multicentric Castleman’s Disease (MCD), indicating their potential contribution to disease progression^43-45^. Secreted viral and host paracrine factors play key roles in the maintenance of the tumor microenvironment and are critical to sustain KSHV-associated malignant tissues. IL-6 and vIL-6 therefore emerge as critical factors during infection. Despite the dependance on these factors, there exists a major barrier to the production of these critical cofactors: the KSHV SOX protein. We have previously demonstrated that IL-6 effectively escape SOX-induced decay but little is known about the regulation of vIL-6. Given the minimal RNA sequence homology between IL-6 and vIL-6, we investigated whether like IL-6, the 3’UTR of vIL-6 alone could confer resistance to SOX. However, we found that the 3’UTR of vIL-6 was also targeted by SOX. Yet, the direct impact of SOX on vIL-6 mRNA remain unclear, as we observed that overall levels of vIL-6 remained high in the KSHV positive cell lines iSLK.219 and BCBL-1. We then showed that vIL-6 nascent RNA levels are significantly increased during lytic reactivation, perhaps as a means to compensate for SOX decay. Indeed, vIL-6 protein levels appeared to remain stable throughout lytic reactivation, suggesting that vIL-6 protein output reflects the delta between increased transcription and increased SOX degradation. This observation suggests distinct regulatory mechanisms governing the stability of viral vs. host transcripts in the face of SOX decay, and it would be interesting to globally assess how individual viral mRNA are affected by host shutoff. One intriguing possibility could be that the expression of vIL-6 and hIL-6 is temporally staggered: early during reactivation, at the onset of SOX-mediated decay, IL-6 predominates as it is unaffected by SOX decay, while vIL-6 transcription ramps up. IL-6 is then destined to be secreted and intracellular level of this cytokine is then replaced by its viral counterpart. This could have extensive effect on the regulation of KSHV infection, with the coordinated effort of IL-6 and vIL-6 to maintain expression of growth factors and inflammatory mediators that support its own viral propagation. Together, the coordinated and sustained expression of both vIL-6 and IL-6 may facilitate long-term viral persistence and contribute to KSHV-associated tumorigenesis.

These observations prompted us to perform RNA-seq analysis in iSLK cells following individual or combined depletion of IL-6 and vIL-6, in order to better define their respective contributions to KSHV-driven gene expression and pathogenesis. Our RNA-seq data revealed a differential impact on gene expression, as expected most of the differentially expressed genes were host-derived rather than viral. This likely reflects the virus dependence on host cellular machinery to support its replication and pathogenesis and suggests that modulation of host pathways plays a central role in the viral life cycle. Notably, host genes involved in signaling and pro-inflammatory responses were significantly affected across all conditions. Interestingly, the pro-inflammatory cytokine IL-1*β* was differentially expressed affected by IL-6 and vIL-6 knock downs: it was downregulated in *Δ*-vIL6 cells but upregulated in si-IL6 samples. This suggests distinct and possibly opposing regulatory roles for IL-6 and vIL-6 in modulating the IL-1*β*-associated inflammatory response during KSHV infection.

Building on these insights, given that vIL-6 and IL-6 activate the JAK/STAT pathway through distinct mechanistic ways, we sought to further investigate their contributions and dynamics during lytic reactivation. As expected, vIL-6 was sufficient to induce the JAK/STAT pathway. In KSHV-infected cells, we observed a similar induction of the pathway but to higher levels of induction at 24 hours when host shutoff is at its peak. This led us to specifically examine the transcription levels of pro-inflammatory factors involved in these pathways. As anticipated, STAT3 was highly expressed, while NF-κB expression remained constantly low compared to STAT3 in KSHV infected cells. Of note, both STAT3 and NF-κB exhibited increased expression at 48h post reactivation, which coincided with the peak expression of vIL-6 as shown in Fig 1B. Notably, STAT3 and NF-κB have been shown to cooperatively promote inflammation, tumorigenesis and disease progression^46-48^. As such, it is not surprising that we observed modulation of expression of STAT3 and NF-κB during lytic infection. Other studies have identified a positive feedback loop involving IL-6, STAT3, NF-κB (known as the IL-6 Amplifier), and potentially other pro-inflammatory factors driving disease and tumor development among other malignancies^48,49^. However, how far this positive feedback loop extends remains unclear. Specifically, during KSHV lytic reactivation, several questions arise: What conditions favor the function of these critical factors in a pro- or anti-viral manner? Are there spatial and temporal constraints for this feedback loop? These are important questions to consider as we continue to explore the complexity of these pathways and their role in disease progression and immunity. Overall, the interactions between vIL-6, IL-6, and the JAK/STAT pathway among others, reveals a complex regulatory network that drives inflammation and tumorigenesis. The potential cooperative activation of STAT3 and NF-κB, along with the crosstalk between other factors from different pathways during lytic infection, highlights their role in KSHV-driven malignancies. While a positive feedback loop involving the combined effect of IL-6 and vIL-6 expression is intriguing, further investigation is needed to uncover the precise factors and conditions that govern their dynamics and the regulation of these pathways. In conclusion, our findings highlight the differential transcriptional regulation of IL-6 and vIL-6, and corroborates that vIL-6 activates host inflammatory signaling through mechanisms distinct from those of IL-6. Consistent with this, our transcriptomic and functional assays show that vIL-6 is sufficient to induce the JAK/STAT pathway and expression of pro-inflammatory factors outside of the context of infection. Notably, during KSHV lytic infection, knockdown of IL-6 further revealed that vIL-6 alone is capable of sustaining JAK/STAT activation and upregulating key pro-inflammatory mediators such as IL6ST, STAT3 and NF-κB. This is particularly important to unravel as one of the only drugs available in the treatment of KSHV-associated lesions like Kaposi Sarcoma is Tocilizumab, a potent antagonist of the IL-6 receptor^50,51^. However, if vIL-6 alone can activate the same pathways as IL-6 yet bypasses the use of the IL-6 receptor, it would be important to design anti-viral treatments that target both IL-6 and vIL-6. Overall, our study helps us better define the contribution of IL-6 and vIL-6 in the pathogenesis of KSHV.

## Materials and Methods

### Cells and transfections

HEK293T cells (ATCC) were grown in Dulbecco’s modified Eagle’s medium (DMEM; Invitrogen) supplemented with 10% fetal bovine serum (FBS). The KSHV-infected renal carcinoma human cell line iSLK.BAC16 (iSLK.WT) and iSLK.219 bearing doxycycline-inducible RTA was grown in DMEM supplemented with 10% FBS. KSHV Lytic reactivation was induced by the addition of 1 μg/ml doxycycline and 1 mM sodium butyrate for 24hr, 48hr or 72hr as indicated. TREX-BCBL-1 cells were cultured in RPMI supplemented with 10% fetal bovine serum (FBS), 2 mM glutamine, 100 U/ml penicillin and 100 mg/ml streptomycin. iSLK.BAC16 vIL-6-STOP and vIL-6-REV (WT) cells (a generous gift from Dr. Yoshihiro Izumiya, UC Davis) were grown in DMEM supplemented with 10% FBS. The vIL-6-STOP virus lacks vIL-6 protein expression due to the addition of stop codons and mutated start codons in the vIL-6 open reading frame within the BAC16 genome. In contrast, the vIL-6 revertant (vIL-6-REV) restores vIL-6 protein expression by reverting the vIL-6-STOP mutations back to the wild-type sequence using BAC recombination^30^. Furthermore, these BAC mutants also incorporated a chicken Bu-1 P2A construct in frame with vIL-6 Wild type or Stop recombinant, enabling the isolation of vIL-6 expressing cells or vIL-6STOP expressing cells by immunoprecipitation using a Bu-1 Mouse anti-chicken, Biotin antibody (#8395-08, Southern Biotech) and Dynabeads MyOne™ Streptavidin C1 beads (#65002, Invitrogen).

For DNA transfections, cells were plated and transfected after 24h when 70% confluent using PolyJet (#504788, SignaGen). For small interfering RNA (siRNA) transfections, cells were reverse transfected using Lipofectamine RNAiMax (#13778075, Thermo) with 10 μM siRNAs. siRNAs were obtained from IDT as Dicer-substrate siRNA (DsiRNA; siRNA IL-6, hs.Ri.IL-6.13.1).

### Plasmids

The vIL-6 3’UTR and coding region were obtained as G-blocks from IDT and cloned into a pcDNA3.1 plasmid downstream of the GFP coding sequence. All cloning steps were performed using in-fusion cloning (#639650, Clonetech-Takara) and were verified by sanger sequencing.

### RT-PCR

Total RNA was harvested using TRIzol according to the manufacture’s protocol. cDNAs were synthesized from 1 μg of total RNA using AMV reverse transcriptase (#M5108, Promega) and used directly for quantitative PCR (qPCR) analysis with the SYBR green qPCR kit (#1725124, Bio-Rad). Signals obtained by qPCR were normalized to 18S unless otherwise noted.

### Immunoblotting

Cell lysates were prepared in lysis buffer (NaCl, 150 mM; Tris, 50 mM; NP-40, 0.5%; dithiothreitol [DTT], 1 mM; and protease inhibitor tablets) and quantified by Bradford assay. Equivalent amounts of each sample were resolved by SDS-PAGE and immunoblotted with each respective antibody in TBST (Tris-buffered saline, 0.1% Tween 20). Antibodies used are Rabbit anti-ORFK2 (ABBIOTEC, 1:200); and Rabbit anti-Vinculin (Invitrogen 1:1000) Rabbit anti-STAT3 (R&D, 1:1000) and Rabbit anti-Cofilin (Invitrogen, 1:2000). Primary antibody incubations were followed by horseradish peroxidase (HRP)-conjugated goat anti-mouse or goat anti-rabbit secondary antibodies (1:5,000; Southern Biotechnology).

### 4-Thiouridine (4sU)

iSLK.219 cells were left latent or lytic reactivated at different timepoints as indicated. Cells were then pulse labeled with DMEM containing 500 μM 4sU (#T4509, Sigma) for 10 minutes, followed by PBS wash and isolation of total RNA with TRIzol. 4sU isolation was performed as described in Garibaldi et al^52^. 4sU isolated RNA was analyzed by RT-qPCR as described above for the indicated genes.

### Enzyme-linked immunosorbent assay (ELISA)

iSLK.219 or TREX-BCBL-1 cells were left latent or lytic reactivated at different timepoints. Cell and supernatants were collected and prepared in lysis buffer (NaCl, 150 mM; Tris, 50 mM; NP-40, 0.5%; dithiothreitol [DTT], 1 mM; and protease inhibitor tablets). IL-6 production was measured using the Invitrogen kit according to the manufacturer’s protocol (#88-7066, Invitrogen).

### Luciferase Assay

HEK293T or iSLK.219 cells were transfected with an ISRE Firefly luciferase reporter vector along with control Renilla luciferase vector from manufacturer (#60613, BPS Bioscience). After, 24 hours, cells were harvested, lysed in Passive Lysis buffer, then luminescence was assessed using the Dual-Luciferase Reporter Assay System (#E2940, Promega) and signal was measured on a luminometer (Tristar 3, Berthold).

### RNA sequencing

vIL-6-STOP or Revertant iSLK cell lines were grown to 80% confluency then transfected with siRNA targeting IL-6 or control scramble siRNA (n=3). Cells were reactivated for 48 hours and immunoprecipitated as previously described, then collected in TRIzol Reagent, and RNA was extracted following the manufacturer’s protocol. Purity of samples were analyzed via bioanalyzer (Agilent 2100). Following poly(A) selection, libraries underwent 75-base paired-end sequencing using the NextSeq500 Mid-150 cycle kit on a NextSeq 500. Read quality was assessed using fastQC. Using Galaxy^53^ reads were aligned to the human genome (hg38) or the KSHV genome (GK18) by HISAT2 and differential expression analysis was performed using FeatureCounts and DESeq2^54,55^. Volcano plots and heatmaps to visualize differential expression were generated using Galaxy.

### Statistical analysis

All results are expressed as means ± standard errors of the means (SEMs) of experiments independently repeated at least three times. Unpaired Student’s t test was used to evaluate the statistical difference between samples. Significance was evaluated with P values as follows: ns., not significant, * p<0.05; ** p<0.01; *** p<0.001.

## Acknowledgments

We would like to thank Dr. Izumiya (UC Davis) for the generous gift of the vIL-6 null KSHV viruses and technical help. The genomics services were performed at the Genomics Resource Laboratory (RRID:SCR_017907), Institute for Applied Life Sciences, University of Massachusetts Amherst, MA.

## Data Availability

The RNA-seq data was deposited in NCBI Gene Expression Omnibus (GEO) database under accession numbers GSE309397.

## Figure Legend

**Supplementary Figure 1: RNA-seq validation**. iSLK vIL-6-STOP and vIL-6-REV cells were treated with siRNA targeting IL-6 or scramble control and left latent or reactivated for 48 hours, then total RNA was harvested followed by RT-qPCR to measure expression of the endogenous genes. (A) CCND1 (B) SOCS3 C) IL6ST. All graphs here are and after display mean ± SEM with individual biological replicates represented as dots. Statistical significance was determined using student *t* test., ns., not significant, * p<0.05, ** p<0.01, *** p<0.001.

**Supplemental Figure 2: Knockdown of IL-6 in iSLK.219 cells increase the expression of vIL-6 and host genes involved in inflammatory signaling**. iSLK.219 cells were treated with siRNA targeting IL-6 (or scramble control) and left latent or reactivated for 24, 48 or 72 hours. Total RNA was harvested and used by RT-qPCR to measure expression of vIL-6 **(A)** or the endogenous expression of IL6ST, STAT3 and NF-kB **(B)**. Graphs display mean ± SEM with individual biological replicates represented as dots. Statistical significance was determined using student *t* test., ns., not significant, * p<0.05, ** p<0.01, *** p<0.001.

