## Supplementary figures and images for "Virally encoded interleukin-6 (vIL-6) coordinates with human IL-6 to modulate cytokine expression during KSHV infection"

### Supplementary Figure 1

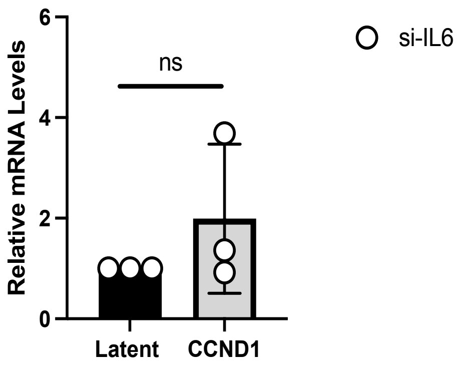

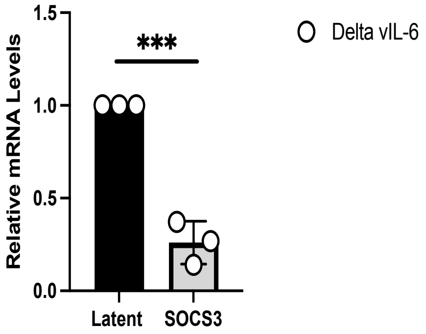

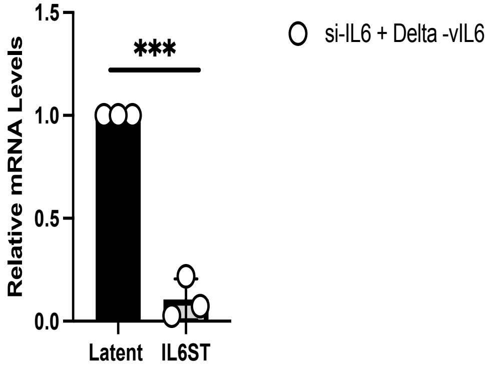

### Supplementary Figure 2

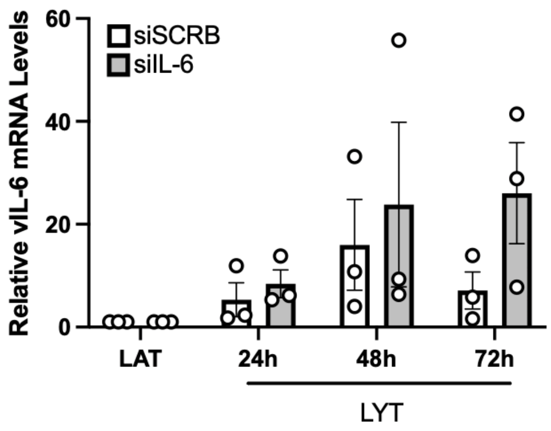

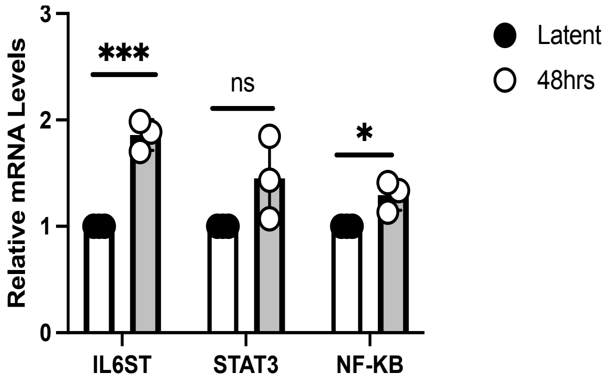
